# A Transcriptional Signature of Induced Neurons Differentiates Virologically Suppressed People Living With HIV from People Without HIV

**DOI:** 10.1101/2024.10.22.619617

**Authors:** Philipp N. Ostermann, Youjun Wu, Scott A. Bowler, Mohammad Adnan Siddiqui, Alberto Herrera, Mega Sidharta, Kiran Ramnarine, Samuel Martínez-Meza, Leslie Ann St. Bernard, Douglas F. Nixon, R. Brad Jones, Masahiro Yamashita, Lishomwa C. Ndhlovu, Ting Zhou, Teresa H. Evering

## Abstract

Neurocognitive impairment is a prevalent and important co-morbidity in virologically suppressed people living with HIV (PLWH), yet the underlying mechanisms remain elusive and treatments lacking. Here, we explored for the first time, use of participant-derived directly induced neurons (iNs) to model neuronal biology and injury in PLWH. iNs retain age-and disease-related features of the donors, providing unique opportunities to reveal novel aspects of neurological disorders. We obtained primary dermal fibroblasts from six virologically suppressed PLWH (range: 27 – 64 years, median: 53); 83% Male; 50% White) and seven matched people without HIV (PWOH) (range: 27 – 66, median: 55); 71% Male; 57% White). iNs were generated using transcription factors NGN2 and ASCL1, and validated by immunocytochemistry and single-cell-RNAseq. Transcriptomic analysis using bulk-RNAseq identified 29 significantly differentially expressed genes between iNs from PLWH and PWOH. Of these, 16 genes were downregulated and 13 upregulated in PLWH iNs. Protein-protein interaction network mapping indicates that iNs from PLWH exhibit differences in extracellular matrix organization and synaptic transmission. *IFI27* was upregulated in iNs from PLWH, which complements independent post-mortem studies demonstrating elevated *IFI27* expression in PLWH-derived brain tissue, indicating that iN generation reconstitutes this pathway. Finally, we observed that expression of the *FOXL2NB-FOXL2-LINC01391* genome locus is reduced in iNs from PLWH and negatively correlates with neurocognitive impairment. Thus, we have identified an iN gene signature of HIV through direct reprogramming of skin fibroblasts into neurons revealing novel mechanisms of neurocognitive impairment in PLWH.

**One sentence summary:** Direct reprogramming of skin fibroblasts into neurons reveals unique gene signatures indicative of HIV infection in the context of viral suppression.

## Introduction

Neurocognitive impairment remains an important co-morbidity of HIV-1 infection in virologically suppressed people living with HIV (PLWH). The introduction of combined antiretroviral therapy (cART) has reduced the prevalence of the most severe forms including HIV-associated dementia. Nonetheless, though milder forms of cognitive impairment are more prominent, the overall burden remains substantial in PLWH with adverse consequences for daily living activities (*1–5*). A 2020 meta-analysis determined the prevalence of HIV-1-related neurocognitive impairment to be 43.9 % (*6*).

The cellular mechanisms responsible for the observed neurocognitive impairment among virologically suppressed PLWH are not well understood but suggested to be multifactorial. It has been shown that HIV-1 can enter the brain as early as two weeks after infection where it infects multiple cell types including T-cells, microglia, brain-resident macrophages, and astrocytes (*7–10*). Neurons are not noticeably infected by HIV-1, yet, the resulting neurotoxic environment impairs neuronal functions driving neurocognitive impairment in PLWH (*10*).

Transcriptomic analysis of post-mortem brain samples derived from PLWH has shown that HIV-1 infection is associated with a differential neural gene expression indicating the involvement of multiple pathways (e.g., axon guidance, endocytosis, synaptic transmission) in the cognitive decline of PLWH

(*11*). However, as Ojeda-Juárez and Kaul recently pointed out, most of these studies lacked suitable non-HIV-1 controls i.e. brain tissue samples derived from people living without HIV (PWOH), which hampers our understanding of direct HIV-1-associated differential neuronal gene expression (*12*). Moreover, the analyzed gene expression may have been affected by different co-morbidities and living situations before death as well as sample preparation after death.

In attempts to overcome these issues, differential gene expression has also been analyzed in neurons from a transgenic HIV-1 gp120 expressing mouse model, which reconstitutes a certain HIV-1-induced neuropathology observed in humans (*13–15*). However, HIV-1 does not naturally infect rodents, cognitive decline in PLWH is influenced by factors beyond gp120, and mouse neuronal biology differs from that of humans. As a result, we are still lacking a neuronal cell system that reflects the multifactorial nature of HIV-1 infection and that allows transcriptional as well as functional analyses of neurons derived from virologically suppressed PLWH.

Recent protocols to generate induced neurons (iNs) by transdifferentiation of participant-derived fibroblasts have made it possible to capture disease-and age-related features of neurons *in vitro* (*16–20*). This made it possible to recapitulate known aspects of neurodegenerative diseases as well as to reveal previously unrecognized underlying disease pathomechanisms in cell culture (*16–18*). Importantly, it has been shown that inducing the pluripotent stem cell state prior to neuronal differentiation of participant-derived cells, as is necessary for the generation of iPSC-derived neurons, erases most age and disease-related characteristics, unlike the iN protocol (*21*). This appears to be particularly important for the study of diseases in which age is implicated in the pathogenesis like Alzheimer’s or HIV-1-related neurocognitive impairment (*17, 22*).

Thus, using a previously published iNs protocol that retains donor-specific disease-and age-related characteristics *in vitro* (*17, 21, 23*), we investigated whether iNs derived from virologically suppressed PLWH show a differential gene expression compared to iNs derived from demographically matched PWOH.

## Results

### Generation of participant-derived induced neurons from people living with or without HIV

We generated iNs from six clinically well-characterized people chronically infected with HIV-1 and virologically suppressed on cART (HIV RNA <50 copies/mL), as well as from seven age-and sex-matched people without HIV-1 (PWOH) as control participants following a recently published protocol (Fig. 1**A**, 1**B**, 1**C**, Table S1) (*21, 23, 24*). PLWH were without neuropsychiatric confounds and underwent comprehensive neurocognitive performance testing (*25*).

**Fig. 1.**
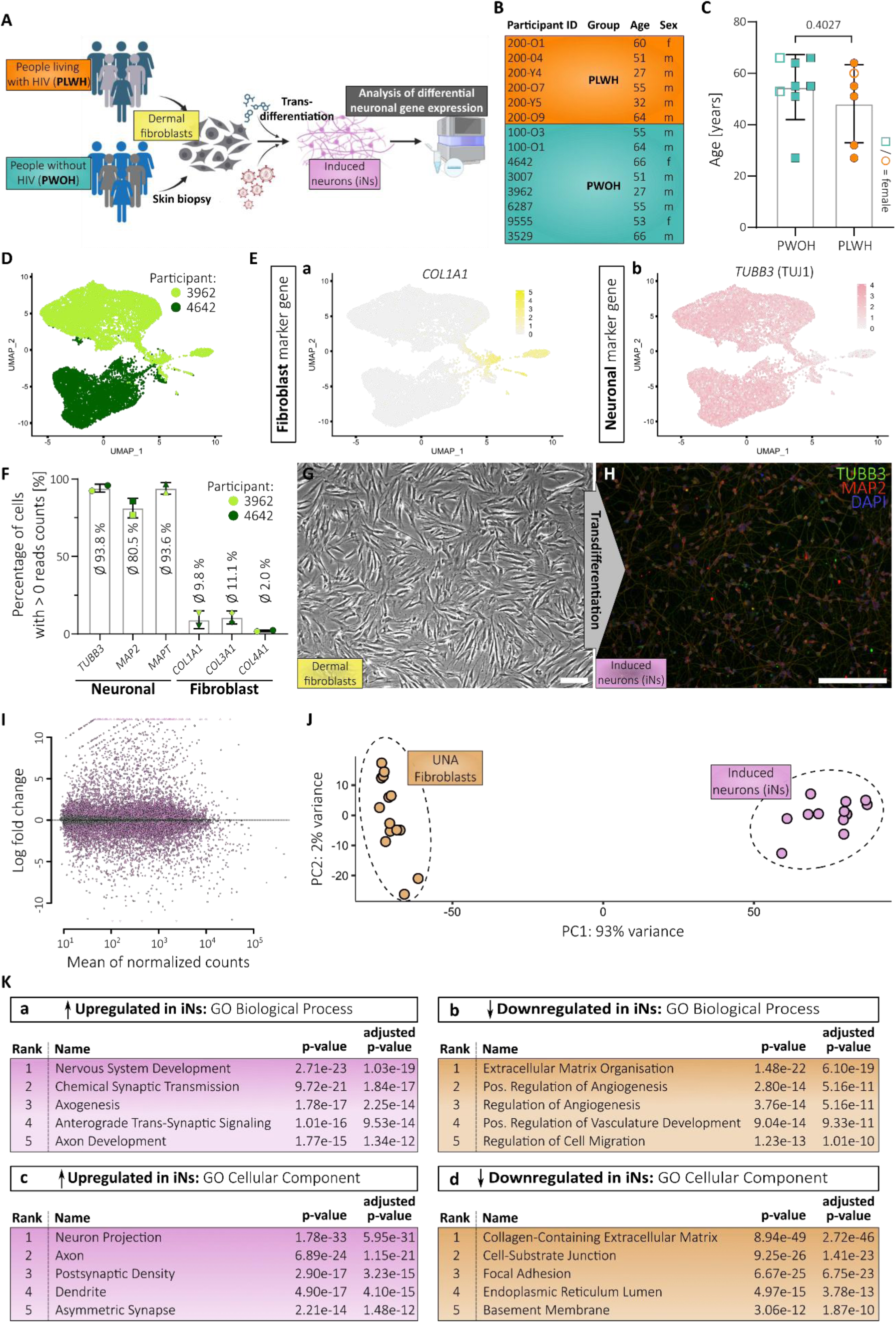
Transdifferentiation of skin fibroblasts derived from people living with HIV and controls generates induced neurons. **(A)** Scheme illustrating study outline. **(B)** Important participant information divided into people living with HIV-1 (PLWH) and without HIV-1 (PWOH). **(C)** Age distribution of the two groups of the study cohort. Statistical significance tested with unpaired, two-tailed *t*-test. Data presented as individual data points with mean ± SD and p-value. **(D)** UMAP plot showing the sample origin of each data point during scRNA analysis. **(E)** Gene expression patterns of fibroblast marker gene *COL1A1* (a) and neuronal marker gene *TUBB3* (b). **(F)** Percentage of cells from scRNA analysis depicted in **(D-E)** that express the annotated neuronal and fibroblast marker genes. Data presented as individual data points with mean ± SD. **(G)** Microscopic image of dermal fibroblasts before transdifferentiation. **(H)** Microscopic image after immunocytochemistry of induced neurons (iNs) 3-days post-FACS and stained for TUBB3 (TUJ1), MAP2 and nuclei (DAPI). Single channel images are provided in Fig. S1F **(G-H)** Scale bars are 20 µm. **(I)** MA plot based on bulk-RNA sequencing showing differential gene expression of all iNs samples compared to UNA fibroblasts. **(J)** PCA plot showing clustering of iNs-and UNA fibroblast-derived RNA samples after bulk-analysis. **(K)** Top 5 ranked gene ontology (GO) terms of Biological Processes (a-b) and Cellular Component (c-d) associated with the significantly up-(a and c) and downregulated genes (b and d) in iNs compared to UNA fibroblasts.

Participant age significantly correlated with the estimated duration of HIV-1 infection in our cohort. The Global Deficit Score (GDS) as measurement for the degree of neurocognitive impairment (NCI) did not correlate with age nor the duration of HIV-1 infection reflecting the fact that multiple, complex factors drive NCI in PLWH (Fig. S1**A**, S1**B**, S1**C**).

To generate iNs, participant-derived dermal fibroblasts were transduced with a lentiviral vector (UNA vector (*23*)) for doxycycline-dependent expression of neuronal transcription factors NGN2 (also NEUROG2), and ASCL1. Transduced fibroblasts were referred to as UNA fibroblasts prior to initiating the transdifferentiation to account for the lentiviral transduction-mediated genetic modification (Fig S1**D**). Treatment of UNA fibroblasts with doxycycline and a cocktail of differentiation factors for 21 days resulted in a mixed population of neurons and non-converted cells as observed by light microscopy and as previously described (Fig S1**E**) (*24*).

To isolate the *bona fide* iNs from this mixed population, we performed fluorescence-activated cell sorting (FACS) targeting polysialylated-neural cell adhesion molecule (PSA-NCAM) on live (DAPI^-^) cells (Fig. S1**D**). To check the purity of the obtained cell population and obtain first insights into the neuronal gene expression, single-cell RNA (scRNA) analysis was conducted with iNs derived from two PWOH (Fig. 1**D**). Gene expression analysis indicated a small subset of cells with a putative fibroblast-associated transcriptome as indicated by expression of *COL1A1*, *COL3A1*, and *COL4A1* that is still contained within the isolated cell population (Fig. 1**E, a**, 1**F**). Nevertheless, the majority of cells expressed neuronal marker genes *TUJ1* (also *TUBB3*), *MAP2*, and *MAPT* (Tau) while lacking fibroblast marker gene expression (Fig. 1**E, b,** 1**F**). Importantly, immunocytochemistry at 3-days post-FACS confirmed TUJ1 and MAP2 protein expression and clearly showed the neuronal morphology of iNs (Fig. 1**G**, 1**H**, S1**F**). Overall, this result was in concordance with a prior study from an independent research group following the same protocol. During their study, a population of approx. 90% TUJ1-positive cells was obtained as analyzed by immunocytochemistry (*17*).

In addition to revealing the expression of typically used pan-neuronal marker genes among our iNs, our scRNA data analysis has confirmed the previous finding of two subpopulations within the iNs: A larger subset of potentially glutamatergic (*SLC17A7*^+^) and smaller subset of potentially GABAergic (*GAD1*^+^) neurons, with virtually no overlap of expression (Fig. S1**G**, S1**H**). Our analysis has further confirmed a lack of choline O-acetyltransferase (*CHAT*) and tryptophan hydroxylase 2 (*TPH2*) expression which would indicate the presence of cholinergic or serotonergic neurons, respectively (Fig. S1**G**).

To expand our validation of the performed transdifferentiation protocol with regard to the entire cohort, we next conducted differential gene expression analysis with bulk-RNA isolated from iNs and their matched UNA fibroblasts derived from all donors (PLWH and PWOH). We harvested RNA while the two matching samples were cultured at the same passage to limit long-term cell culture-mediated effects between the UNA fibroblasts and their corresponding iNs. As a result, the only difference between the UNA fibroblasts and their matched iNs has been the 21 days of transdifferentiation and subsequent cell sorting. The differential gene expression analysis showed that the morphological transition into the neuronal phenotype was accompanied by drastic transcriptional changes with over 10,000 genes being differentially expressed in the iNs compared to the UNA fibroblasts (p-adj. < 0.05, log2fc > +/-0.5) (Fig. 1**I**, S2). This finding is supported by principal component analysis (PCA), which depicted a clear separation of the clustered iN and UNA fibroblast samples (Fig. 1**J**).

The two fibroblast cultures derived from PWOH participants 100-O1 and 100-O3 were generated in our laboratories together with the PLWH fibroblast cultures following in-house protocols (Table S1). Importantly, the respective UNA fibroblast samples derived from participant 100-O1 and 100-O3 showed no association with the PLWH samples that were likewise generated but instead clustered with the rest of the PWOH samples that were obtained elsewhere (Fig. S1**I**). This indicated that the method of fibroblast culture generation did not affect our downstream analysis and corroborated the validity of our samples for the described analyses.

To verify the neuronal biological state of the iNs, gene set enrichment analysis was performed determining Gene ontology (GO) terms associated with UNA fibroblast to iN transdifferentiation (Fig. 1**K**). As expected, GO terms of biological processes associated with the upregulated genes after transdifferentiation were linked to neuronal development and function (Fig. 1**K**, **a**). This was corroborated by the top GO terms of cellular components, which corresponded to neuron-specific compartments (Fig. 1**K**, **c**). Further, downregulated genes following transdifferentiation into iNs were associated with GO terms corresponding to fibroblasts function and related cellular compartments, respectively, underlining the loss of fibroblast-associated characteristics over the 21 days of transdifferentiation (Fig. 1**K**, **b** -**d**). Together, the gene set enrichment analysis on bulk-RNA from all donors strongly support the transdifferentiation of our participant-derived fibroblasts into neurons.

In summary, these analyses demonstrate successful execution of the previously published protocol for the generation of participant-derived iNs from our cohort of six PLWH and seven matched controls of PWOH. Hence, we obtained neurons via a protocol, which preserves age-and disease state-associated biological changes and that allows us to now reveal biologically plausible neuronal differences between virologically suppressed PLWH and PWOH.

### PLWH-derived iNs exhibit statistically significant differentially expressed genes compared to iNs from PWOH

After confirming our ability to successfully transdifferentiate participant-derived fibroblasts from PLWH and PWOH into iNs, we aimed to investigate if the presence of chronic HIV-1 infection in the context of concurrent treatment with combination antiretroviral therapy (cART) affects the gene expression signature of the iNs. For this, we used the obtained bulk-RNA sequencing data to compare gene expression profiles of the iNs derived from PLWH to those derived from PWOH. This transcriptome analysis identified 29 differentially expressed genes (DEGs) between PLWH-and PWOH-derived iNs (p-adj. < 0.05, log2fc > +/-0.5) (Fig. 2**A**, 2**B**, Table S2). Of these, 13 genes were upregulated, and 16 genes downregulated in the iNs derived from PLWH (Fig. 2**B**). Six of these 29 genes were likewise differentially expressed between the matched UNA fibroblast samples of PLWH vs. PWOH indicating a broader, perhaps cell-type independent effect in PLWH on these genes (Fig. 2**C**).

**Fig. 2.**
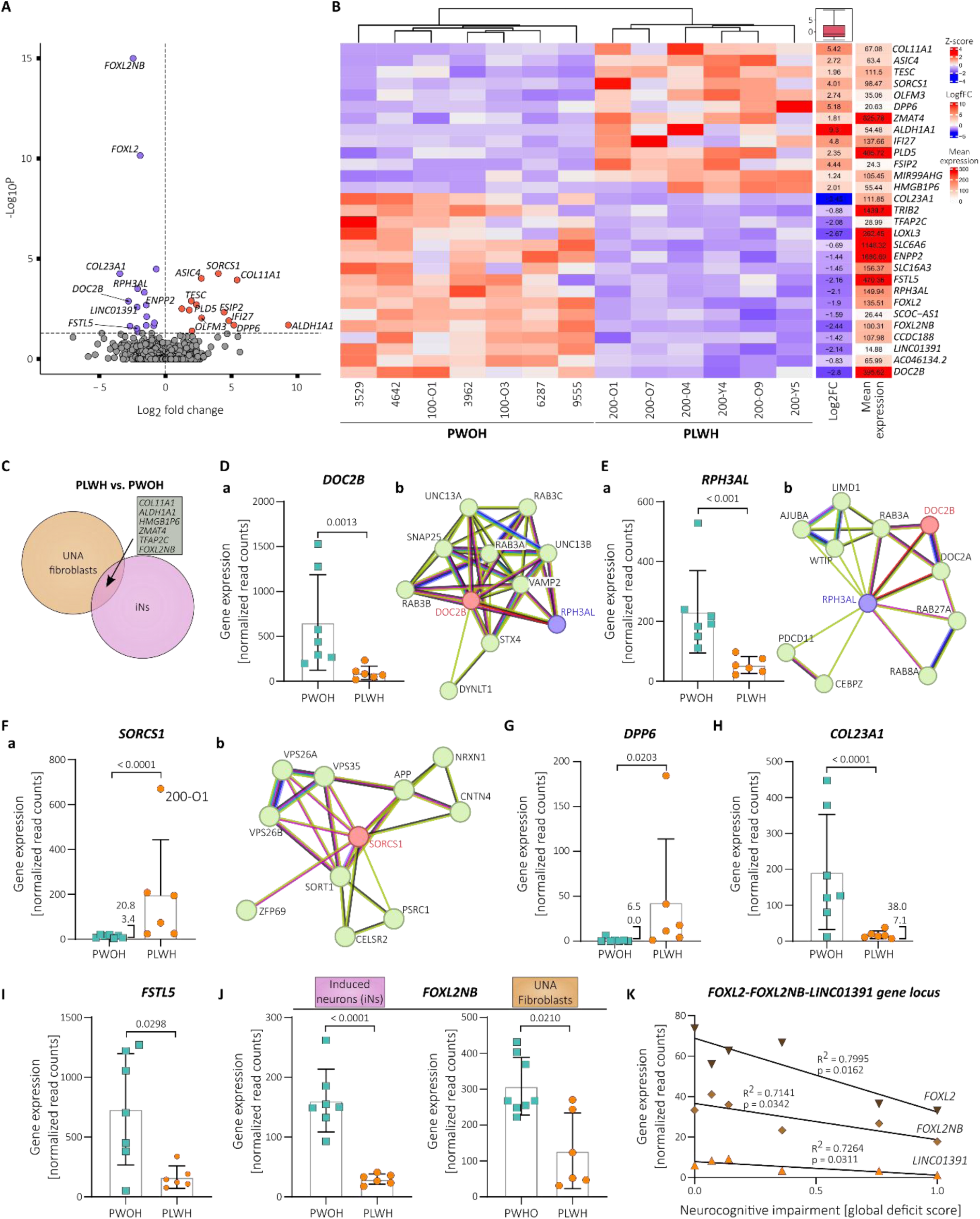
PLWH-derived iNs exhibit statistically significant differentially expressed genes compared to iNs from PWOH. **(A)** Volcano plot showing the 29 statistically significant (p-adj. < 0.05, log2fc > +/- 0.5) differentially expressed genes (DEGs) in PLWH-derived iNs compared to PWOH-derived iNs following bulk-RNA analysis iNs. Y-axis plots the p-adj. values and the dotted line indicates the selected cut-off of p-adj. < 0.05. **(B)** Heatmap showing the clustering of PLWH-vs. PWOH-derived iNs RNA samples based on expression levels of the 29 DEGs while displaying the log2 fold change and mean expression. **(C)** Venn diagram showing the numbers of DEGs between the PLWH-and PWOH-derived samples for the UNA fibroblasts and iNs. The gene symbols of the six genes that are differentially expressed in both cell types are displayed in the grey box. **(D-F)** Gene expression levels in PLWH-vs. PWOH-derived iNs (a) and STRING protein association networks (b) of *DOC2B* **(D)**, *RPH3AL* **(E)**, and *SORCS1* **(F)**(*33, 34*). **(G-I)** Gene expression levels in PLWH-vs. PWOH-derived iNs of *DPP6* **(G)**, *COL23A1* **(H)**, and *FSTL5* **(I)**. **(J)** Gene expression levels in PLWH-vs. PWOH-derived iNs and UNA fibroblasts of *FOXL2NB*. **(D-J)**. Data presented as individual data points with mean ± SD and p-adj-value derived from the conducted Wald test corrected for false discovery rates (FDR) using the Benjamini-Hochberg method (see methods section). **(K)** Correlation of gene expression levels of *FOXL2*, *FOXL2NB*, and *LINC01391* with neurocognitive impairment in the PLWH study group (n =6). Data points are individual values and lines depict linear regression functions.

We considered the remaining 23 DEGs to be iN-specific in our setting. This list of iN-specific DEGs contained several genes including *DOC2B, RPH3AL, SORCS1,* and *DPP6* that are associated with either known or presumed neuronal functions as well as dysfunctions (*26–30*). These candidates may present novel pathways relating to neuronal function in virologically suppressed PLWH.

### Differential expression of genes related to synaptic transmission pathways in PLWH iNs compared to PWOH iNs

Expression of the iN-specific DEG double C2 domain beta (*DOC2B*) was reduced by 6.95-fold in PLWH compared to PWOH on average (Fig. 2**D**, **a**). *DOC2B* is readily expressed in the human brain and important for neuronal activity (*31*). DOC2B has been identified as a cytosolic Ca^2+^ sensor that mediates spontaneous neurotransmitter release (*26*). It was shown that a double knockdown of the double C2 domain proteins DOC2A and DOC2B results in a decrease in spontaneous transmitter release from hippocampal neurons, which could be rescued by expressing DOC2B (*26*). A subsequent study later described its role in hippocampal synaptic plasticity (*32*). DOC2B interacts with several proteins that are important for neurotransmitter release including components of the SNARE complex as shown by pulldown analysis (*26*) as well as revealed by STRING network analysis (Fig. 2**D**, **b**) (*33, 34*). Interestingly, the resulting graph of the network analysis also depicts a link to rabphilin 3A like (without C2 domains) (*RPH3AL*, previously *Noc2*), another gene that was significantly downregulated in the PLWH-compared to the PWOH-derived iNs (Fig. 2**E**, **a** - **b**). This co-reduction may be due to shared gene regulatory elements that supposedly control *DOC2B* and *RPH3AL* transcription on chromosome 17 (*35, 36*). RPH3AL is a Rab effector protein associated with release of secretory vesicles (*27*). It is expressed presumably at low levels in the brain (*28*), and a recent GWAS study showed that the *RPH3AL* missense mutation rs117190076 increases the risk for late-onset Alzheimer’s disease (*37*). Hence, downregulation of the genomic locus containing *DOC2B* as well as *RPH3AL* may affect synaptic vesicle release in two different ways.

In contrast to *DOC2B* and *RPH3AL*, olfactomedin 3 (*OLFM3*) showed increased expression in iNs derived from PLWH compared to PWOH (Fig. 2**A**, 2**B**). OLFM3 protein is expressed in neurons of the cortex, and hippocampus (*38*). It has been shown to bind different subunits of the postsynaptic AMPA receptor (α-amino-3-hydroxy-5-methyl-4-isoxazole-propionic acid receptor), namely GRIA1, and GRIA2 and its overexpression in the mouse hippocampus affects their membrane expression (*38*). In this context, increased OLFM3 expression has been linked to epilepsy because it was suggested to alter AMPA receptor activity (*38*).

We conclude that iNs derived from PLWH exhibit differential expression of genes directly involved in pre-as well as postsynaptic processes when compared to their counterparts with PWOH.

### Expression of Alzheimer’s disease-associated *SORCS1* is increased in PLWH-derived iNs

An important pathway in the development of Alzheimer’s disease is the dysregulated processing of amyloid precursor protein (APP), which leads to intracellular accumulation of Aβ. In addition to apolipoprotein E (ApoE), amyloid-beta (Aβ), and Tau protein (hTau) (*39, 40*), sortilin related VPS10 domain containing receptor 1 (*SORCS1*) expression is amongst the best known risk factors in Alzheimer’s disease. SORCS1 is associated with different components of the Aβ pathway and has been specifically shown to play a role in aberrant APP trafficking (Fig. 2**F**, **b**) (*29, 41*). In addition to its role in Alzheimer’s, SORCS1 is a general regulator for intracellular trafficking and is important for maintaining neuronal functionality by, for instance, sorting the AMPA glutamate receptor (AMPAR) and synaptic adhesion molecule neurexin (NRXN), which ensures proper glutamatergic transmission (*42*). We observed significantly elevated *SORCS1* expression levels in the iNs of PLWH when compared to the PWOH iNs in our study (Fig. 2**F**, **a**). To our knowledge, this is the first reported association of *SORCS1* gene expression with HIV-1 infection in the published literature.

Interestingly, the association between *SORCS1* SNPs and Alzheimer’s disease appears to exhibit a sexual dimorphism, with a stronger correlation observed in women (*40, 43*). In our cohort of PLWH, *SORCS1* expression was found to be 3-fold higher in the iNs from the participant of female sex (200-O1) compared the iNs from the participant with the second highest *SORCS1* expression (Fig. 2**F**, **a**). However, the limited sample size prevents us from drawing conclusions regarding the effect of sex on *SORCS1* expression.

### Expression of the potassium ion channel auxiliary factor DPP6 is increased in iNs derived from PLWH

Dipeptidyl peptidase like 6 (*DPP6,* previously *DPPX*) is another iN-specific DEG with a known role in neuronal function. *DPP6* exhibited increased expression levels in the iNs derived from PLWH compared to PWOH (Fig. 2**G**). DPP6 RNA and protein expression throughout the human body is predominantly found within the brain with low region specificity (available from v23.0proteinatlas.org; https://www.proteinatlas.org/ENSG00000130226-DPP6) (*44, 45*). It is an important auxiliary factor of potassium ion (K^+^) channels, and its expression is associated with synaptic function and impairments in learning and memory (*30, 46–48*). In addition, the NHGRI-EBI GWAS catalog (*49*) includes *DPP6* SNPs associated with cognitive decline (GCST009443; GCST90308745) (*50, 51*), and hippocampal volume (GCST90104700) (*52*). Despite obtaining a low average expression of *DPP6* in our assay (Fig. 2**G**), it is notable that the performed bulk-RNA sequencing resulted in zero reads for *DPP6* RNA in 4 out of 7 iNs samples derived from PWOH whereas *DPP6* expression was detected in all PLWH iNs samples (Fig. 2**G**). Studies investigating DPP6 function in the past have typically used DPP6 knockdown or knockout experiments, which enables only the analysis of lack of DPP6 expression (*46–48*). Thus, it is difficult to draw any conclusions based on their results about a potential pathophysiological effect of increased *DPP6* levels as it is the case in the PLWH iNs. However, a single study on schizophrenia that generated participant-derived induced pluripotent stem cell (iPSC)-derived neurons found increased *DPP6* transcript levels in neurons from schizophrenia patients compared to healthy controls (*53*). In that study, multi-electrode-array recordings and calcium imaging demonstrated decreased neuronal activity in the neuronal cultures of schizophrenia patient-derived cells. Moreover, shRNA-mediated reduction of *DPP6* levels as well as pharmacological inhibition of the K^+^ channel Kv4.2 reversed this observed hypoexcitability, and hypoactivity suggesting a causal relationship between increased *DPP6* levels and decreased neuronal activity. We therefore hypothesize that the iNs of PLWH, which showed increased *DPP6* expression compared to matched controls, may exhibit the same aberrant excitability -a link that warrants further investigation.

### Differential expression of extracellular matrix-associated genes in iNs from PLWH compared to those from PWOH

The extracellular matrix (ECM) plays an important role in neuronal development, and function (*54*) and the dysregulation of ECM-associated proteins including members of the collagen family has been linked to neurodegeneration (*55*). Notably, neurons and non-neuronal glial cells actively shape their surroundings by expressing and secreting various ECM proteins, which is required for important processes, e.g., synaptic plasticity (*54, 55*).

In this study, iNs derived from PLWH exhibited significantly differential gene expression of several ECM-associated proteins when compared to iNs derived from PWOH. In the majority of cases, similar changes in gene expression patterns were not noted in the matched UNA fibroblasts samples indicating neuron-specific effects.

Collagen type XXIII alpha 1 chain (COL23A1) is an ECM-associated protein that may reveal mechanic insights into HIV-1-related neurocognitive decline. Its expression was more than 8-fold reduced in iNs derived from PLWH compared to PWOH (Fig. 2**H**). Further, we obtained raw read counts for *COL23A1* RNA via bulk-RNA sequencing from only four UNA fibroblast samples ranging from 1 -12 raw reads, which substantiates the iN-specific *COL23A1* expression in our assay (data not shown). Indeed, *COL23A1* expression is found across the human brain in multiple cell types including neurons, and astrocytes (available from v23.0proteinatlas.org; https://www.proteinatlas.org/ENSG00000050767-COL23A1/brain) (*44, 45*). COL23A1 is a type II membrane protein belonging to the transmembrane collagen family (*56*) and despite a general lack of knowledge concerning its role in neural functions, SNPs of its gene are associated with the rate of cognitive decline in Alzheimer’s disease (GCST010567) (*57*) and memory performance (GCST90104696) (*52*).

In addition to *COL23A1*, PLWH-derived iNs exhibited differential expression of the collagen family member *COL11A1* (collagen type XI alpha 1 chain) (Fig. 2**B**). While potentially of interest, unlike findings with *COL23A1*, the increased expression of COL11A1 in PLWH samples when compared to PWOH controls was not restricted to iNs and was also observed in the UNA fibroblasts (Fig. 2**C**). Of note, *COL11A1* is expressed in the brain as well as the skin (available from v23.0proteinatlas.org; https://www.proteinatlas.org/ENSG00000060718-COL11A1) (*44, 45*).

Besides structural proteins like the collagen family members, different secreted proteins are also associated with the ECM. Our iNs from PLWH showed decreased expression levels of *ENPP2* (Fig. 2**B**). The *ENPP2* gene encodes ectonucleotide pyrophosphatase/phosphodiesterase 2, which is better known as autotaxin. Autotaxin is an enzyme secreted into the ECM by different cell types throughout the human body including neural cells (*58*). Autotaxin exerts biological functions by processing lysophosphatidylcholine into lysophosphatidic acid (LPA), which then binds to one of its several G-protein coupled receptors (LPA1-LPA6) (*58*). Interestingly, LPA signalling is involved in numerous physiological processes including neurogenesis, neuronal differentiation, synapse formation, migration, and cortical development (reviewed in (*58*)). Hence, decreased expression of the *ENPP2* gene in neural cells may affect brain functioning via dysregulated LPA signalling.

Follistatin like 5 is also a secreted protein, which is expressed throughout the human brain, predominantly in the cerebellum, in inhibitory as well as excitatory neurons (available from v23.0proteinatlas.org; https://www.proteinatlas.org/ENSG00000168843-FSTL5) (*44, 45*). Although its role in physiological brain processes is not well described, single nucleotide polymorphisms (SNPs) found within the *FSTL5* gene are associated with general cognitive ability (GCST006269) (*59*), dementia and Alzheimer’s disease in non-APOE ε4 allele carriers (GCST90244035; GCST90244033) (*60*), and working memory (GCST006930) (*61*) as annotated in the NHGRI-EBI Catalog of human genome-wide association studies (GWAS) (downloaded 10/03/2024) (*49*) . We found that expression levels of *FSTL5* were on average 4.4-fold decreased in PLWH-derived iNs compared to PWOH iNs (Fig. 2**B**, 2**I**).

### Protein-protein interaction network mapping supports differential ECM organization, and synaptic transmission in PLWH iNs and indicates neuronal apoptosis as another affected pathway

Since the dysregulation of a single gene product likely affects several other gene products, protein-protein-interaction (PPI) network mapping has become a powerful tool to analyze complex pathomechanisms. Furthermore, different PPI databases (e.g, IntAct, HuRI) that allow network mapping based on curated experimental data sets have made it possible to bridge the gap between RNA expression and protein interactions without the necessity for additional co-precipitation or proximity ligation assays (*62, 63*).

After identifying 29 DEGs that distinguish PLWH-and PWOH-derived iNs by transcriptomic analysis, we sought to investigate whether PPI network mapping reveals insights into the affected cellular pathways. To this end, we determined the 1^st^-order interaction partners for the gene products of the 29 DEGs between PLWH-and PWOH-derived iNs using the IntAct Molecular Interaction Database (*62*) and The Human Reference Interactome (HuRI) (Fig. S3**A**, Table S3).

Gene set enrichment analysis revealed that biological processes associated with the resulting PPI network were related to neuronal apoptosis (Fig. S3**B**, **a**, Table S4). Further, this analysis supported the idea that the organization of the ECM is affected in iNs derived from PLWH compared to PWOH (Fig. S3**B**, **b**). In addition to the ECM, the PPI network analysis substantiated an influence of the 29 DEG on synaptic transmission (Fig. S3**B**, a and c).

When analyzing the obtained PPI network with regard to associated diseases, it was therefore not surprising to find associations with terms like neurodegenerative disease, dementia, or Alzheimer’s disease (Fig. S3**B**, d – e) (*64, 65*).

We conclude that the DEGs in iNs from virologically suppressed PLWH may drive various disease-associated pathways in the CNS by direct protein-protein interactions of the respective gene products.

### Expression of the *FOXL2NB*-*FOXL2*-*LINC01391* genome locus is reduced in PLWH-derived iNs and associated with the degree of neurocognitive impairment

Expression of the FOXL2 neighbour gene (*FOXL2NB*, previously *C3orf72*) is significantly downregulated in PLWH-when compared to PWOH-derived iNs but also in the UNA fibroblasts (Fig. 2**C**). However, we found this effect to be more pronounced in the iN samples (Fig. 2**J**). This suggests that the differential *FOXL2NB* expression between PLWH and PWOH in the iN samples is not an experimental artifact mediated by the choice of our original cell type, the participant-derived primary dermal fibroblasts, but rather a cell type-independent effect that appears to be more prominent in neurons than in fibroblasts.

Furthermore, expression levels of the transcription factor forkhead box L2 gene (*FOXL2*), and the long non-coding RNA LINC01391 were significantly reduced only in the PLWH iNs samples and not the PLWH UNA fibroblasts (Fig. 2**B**, 2**C**). We found this of particular interest because the three genes, *FOXL2NB*, *FOXL2*, and *LINC01391* are located in close proximity to each other on human chromosome 3 and their expression is controlled by shared gene regulatory elements as annotated in the *GeneHancer* database (Fig. S4) (*35, 36*). Thus, the transcription rate at this genomic locus may be decreased in neurons of PLWH. Interestingly, little is known about the function of LINC01391, and FOXL2NB in the brain but *FOXL2* has been very recently associated with Alzheimer’s disease (*66*). Kavoosi et al. have used published microarray expression data on tissue from different brain regions (e.g., frontal, temporal, and entorhinal cortex) obtained from healthy controls, asymptomatic and symptomatic Alzheimer’s patients to identify *FOXL2* via microRNA-mRNA regulatory networks (*66, 67*).

Moreover, expression levels of *FOXL2NB*, *FOXL2*, and *LINC01391* showed a negative correlation with the global deficit score in our cohort, i.e., the more pronounced the NCI the lower their expression levels (Fig. 2**K**). Overall, this data set may indicate a novel pathway in neurocognitive decline among PLWH and give rise to novel marker genes for future experimental studies.

### Autopsy-tissue samples from the NeuroAIDS Tissue Consortium confirm increased levels of the iN-specific DEG *IFI27* in the brains of PLWH

Sustained inflammation of the CNS is considered a major factor in the development of HIV-1-related neurocognitive impairment. Several genes of the inflammatory signaling cascade have been identified so far that may contribute to this including *ISG15*, *IFIT1*, *IF44*, and *IFITM1* (*11, 68–70*).

Interestingly, our differential gene expression analysis revealed interferon alpha-inducible protein 27 (*IFI27*) to be upregulated in iNs derived from virologically suppressed PLWH when compared to PWOH (Fig. 3**A**, **a**). This *IFI27* upregulation was not observed in the UNA fibroblasts and was therefore suggested to be iN-specific (data not shown). STRING network analysis clearly showed that *IFI27* is closely associated with *ISG15*, *IFIT1*, *IF44*, *IFITM1*, and many additional genes of the inflammatory pathway (Fig. 3**B**). Hence, we concluded that based on our approach to investigate differential neuronal gene expression in virologically suppressed PLWH by generating iNs, that *IFI27* may also play a role in HIV-1-associated neuroinflammation.

**Fig. 3.**
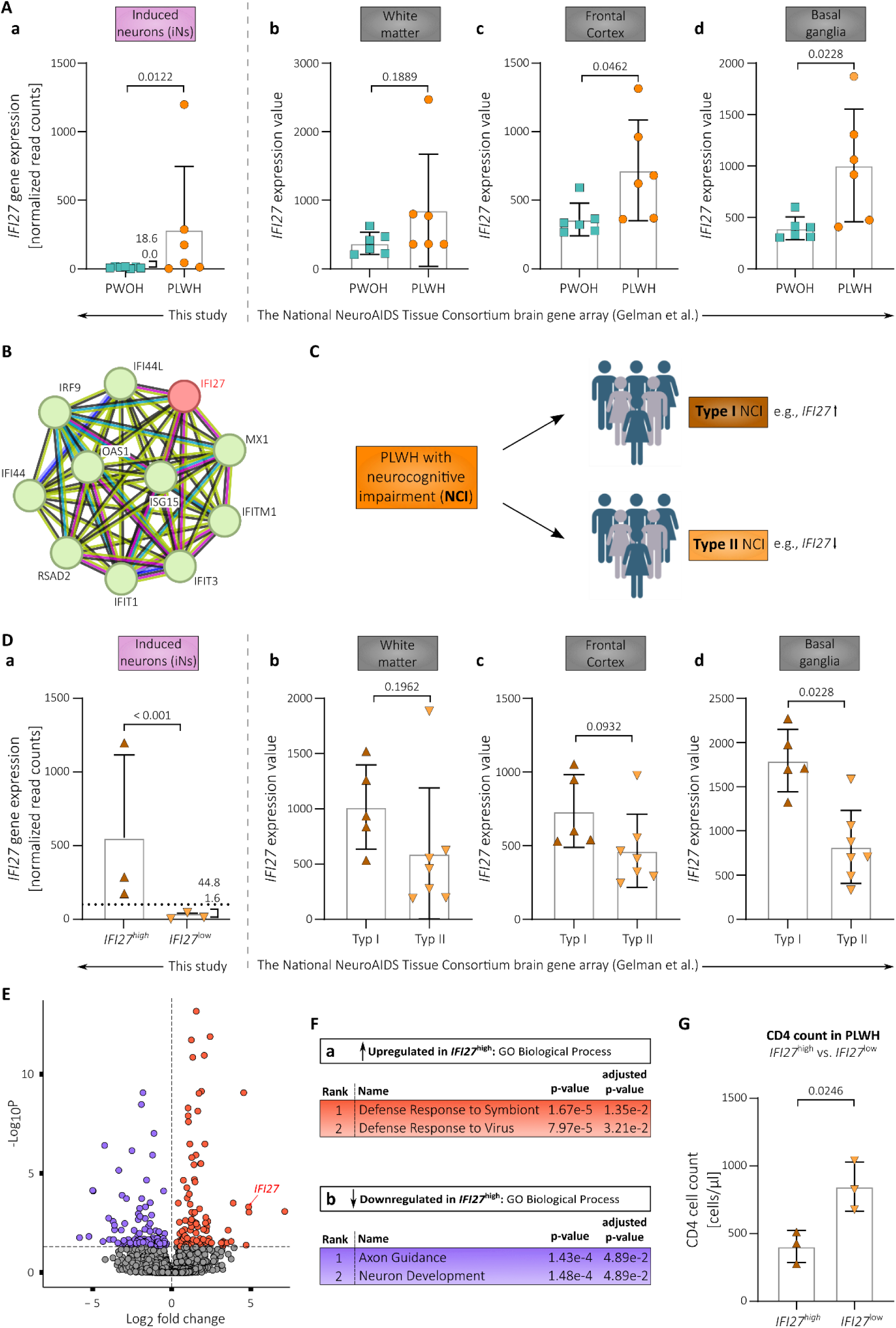
*IFI27* expression levels are increased in PLWH-derived iNs and post-mortem brain tissue samples compared to PWOH-derived samples. **(A)** *IFI27* gene expression levels in PLWH-vs. PWOH-derived iNs (a) and post-mortem brain tissue samples (b-d). **(B)** STRING protein association network of *IFI27*(*33, 34*). **(C)** Scheme illustrating the concept of type I and II neurocognitive impairment in PLWH according to Gelman et al. together with the associated *IFI27* expression (*11*) (D) *IFI27* gene expression levels in PLWH *IFI27*^high^ vs. *IFI27*^low^ iNs (a) and type I vs. type II PLWH-derived post-mortem brain tissue samples (b-d). **(E)** Volcano plot showing the statistically significant (p-adj. < 0.05, log2fc > +/-0.5) differentially expressed genes (DEGs) in PLWH *IFI27*^high^ vs. *IFI27*^low^ iNs based on our bulk-RNA analysis. **(F)** Top 2 ranked gene ontology (GO) terms of Biological Processes associated with the significantly up-(a) and downregulated genes (b) in PLWH *IFI27*^high^ vs. *IFI27*^low^ iNs. **(G)** CD4^+^ T cell counts in PLWH divided into *IFI27*^high^ vs. *IFI27*^low^ participants. Statistical significance tested with unpaired, two-tailed *t*-test **(A** b-d**, D** b-d**, G)** or derived from the conducted Wald test corrected for false discovery rates (FDR) using the Benjamini-Hochberg method on whole-transcriptome data **(A** a, **D** a). Data presented as individual data points with mean ± SD. *IFI27* expression values in post-mortem brain tissue samples derived from Gelman et al.(*11*).

Given this strong connection to the inflammatory signals that have been previously linked to HIV-1-related neurocognitive impairment outlined above (Fig. 3**B**), we mined the literature to find pre-existing evidence supporting our conclusion that *IFI27* may be involved as well. Importantly, we found three independent studies comparing gene expression in post-mortem brain samples between PLWH and PWOH that found significantly increased *IFI27* expression in brain tissue derived from PLWH (*11, 69, 71*).

Solomon et al. compared gene expression in frontal white matter tissue between 34 PLWH (≥ 45 years old) on cART and 24 age-matched PLWH via gene expression profiling and showed a more than 2-fold increase in *IFI27* expression in the PLWH-derived brain tissue samples (*69*).

Gabuzda et al. very recently likewise performed gene expression profiling and found an 1.73-fold increased *IFI27* expression in the frontal lobe white matter samples derived from 28 PLWH (≥ 40 years old) on cART when compared to samples derived from 20 age-and sex-matched PWOH (*71*).

Lastly, Gelman et al. performed a gene expression array with post-mortem tissue from the frontal cortex (neocortex), white matter, and basal ganglia (neostriatum) obtained from the National NeuroAIDS Tissue Consortium in 2012 (*11*). By analyzing their publicly available gene array data set on post-mortem brain tissue, we observed that *IFI27* expression levels are significantly elevated in the basal ganglia, and frontal cortex of PLWH without NCI when compared to PWOH (Fig. 3**A**, **c** **-d**). The increase in *IFI27* expression observed for the analyzed white matter samples was not significant but nevertheless showed a trend towards upregulation (Fig. 3**A**, **b**). Together, this shows that *IFI27* expression is increased in the brains of PLWH and that our here generated iNs derived from virologically suppressed PLWH reconstitute this upregulation.

Based on a comparison of DEGs between PWOH and PLWH with NCI that either developed HIV encephalitis (HIVE) or not, Gelman and colleagues have suggested the presence of two distinct pathomechanisms that lead to NCI in PLWH irrespective of cART: With inflammatory changes (Type I NCI) and without inflammatory changes (Type II NCI) (Fig. 3**C**). Type I and II NCI in PLWH supposedly underlie different biological pathways and several marker genes including *IFI27* were identified whose expression levels together could distinguish between the two types (*11, 72*).

In this regard, expression of *IFI27* alone may not be enough to reliably distinguish between the two types because its expression differed significantly only in the basal ganglia samples (Fig 3**D**, **b -c**). However, to nevertheless test whether the distinction of type I and II NCI may be preserved after transdifferentiation of dermal fibroblasts into iNs, we first divided our PLWH study group into *IFI27* low and high expressing participants (*IFI27*^low^ vs. *IFI27*^high^) (Fig. 3**D**, **a**). Based on the average foldchange between *IFI27* expression in PWOH vs. PLWH with type I NCI in the post-mortem samples of the basal ganglia (∼ 5-fold), we chose a cut-off that was 5-fold the average *IFI27* expression in iNs derived from PWOH (= 51.50 normalized read counts).

Next, we performed differential gene expression analysis to compare the transcriptional profile of the *IFI27*^high^ with the *IFI27*^low^ PLWH iNs, the first now serving as putative surrogate model for type I NCI (*IFI27*^high^), and the latter for type II NCI neurons (*IFI27*^low^). We identified 215 DEGs (p-adj. < 0.05, log2fc > +/- 0.5), of which 106 were downregulated and 109 upregulated in the *IFI27*^high^ PLWH iNs (Fig. 3**E**, S5**A** Table S5). The performed PCA did not reveal any obvious clustering between the two groups but on the other hand did not provide evidence against it (Fig. S5**B**). This might have been due to the low sample size in this context.

Gene set enrichment analysis showed that a significant number of the upregulated genes were associated with antiviral defense (Fig. 3**F**, **a**). This is in line with the Ingenuity Pathway Analysis performed by Gelman and colleagues, who found canonical pathways of antiviral defense mechanisms upregulated in their type I NCI samples as well (*11*). Importantly, GO terms associated with the downregulated genes in the *IFI27*^high^ PLWH iNs were related to neuronal development and the formation of neuronal processes with the GO term ranked as number 1 being *Axon Guidance*, a term that has been likewise found to be associated with the downregulated genes in type I NCI samples in the study by Gelman et al. (Fig. 3**F**, **b**) (*11*). Hence, our analysis indicated that studying biological pathways underlying the distinction of type I and II NCI may be possible by transdifferentiation of participant-derived dermal fibroblasts into iNs.

Lastly, to investigate whether the made distinction between *IFI27*^high^ and *IFI27*^low^ PLWH iNs can be associated with any biological parameters indicative of disease state or donor characteristics, we analyzed possible associations with the age, duration of HIV-1 infection, neurocognitive impairment, and CD4^+^ cell count of the respective PLWH donors. Here, we observed a significant association of the CD4^+^ T-cell count at the day of the skin bunch biopsy on *IFI27* expression in the iNs (Fig. 3**G**, S5**C**, S5**D**, S5**E**).

We conclude that post-mortem sampling of brain tissue from PLWH confirms that the elevated *IFI27* expression levels we observed in the here generated PLWH-derived iNs also occur in the brains of PLWH. Further, the identified DEGs between *IFI27*^high^ and *IFI27*^low^ PLWH iNs indicate that the distinct mechanisms responsible for type I and II NCI are conserved in iNs and thus support iNs as a novel model system to study cognitive decline in PLWH.

Overall, differential gene expression analysis on participant-derived iNs revealed 29 DEGs between PLWH-and PWOH-derived iNs potentially revealing novel mechanisms and supporting previous concepts of HIV-1-related neuroinflammation and neurocognitive decline.

## Discussion

In this study, using a recently described protocol to generate iNs that retains donor-specific disease-and age-related characteristics *in vitro*, we set out to investigate whether iNs derived from virologically suppressed PLWH exhibit differential gene expression compared to iNs from matched PWOH (*24*). We identified an iN gene signature of HIV-1 comprising 29 DEGs including genes associated with neuronal functions implicated in cognitive decline.

The performed transdifferentiation to generate iNs resulted in around 90 % cells expressing pan-neuronal marker gene *TUBB3* (TUJ1). This is in line with the rate of iNs obtained by the research group that originally established this protocol and applied it to the study of Alzheimer’s disease (*17*). Further, the same protocol resulted in about 15 - 20% cells expressing the glutamatergic neuronal marker *SLC17A7* and about 5 % cells expressing the GABAergic neuronal marker *GAD1* in a previous study (*16*). Consistently, we have observed rates of 32.8 % *SLC17A7^+^* and 7.7 % *GAD1^+^* cells among our iNs underlining the feasibility and comparability of the published protocol among different labs.

Subsequent transcriptomic analysis of the participant-derived iNs revealed 29 DEGs between virologically suppressed PLWH and PWOH. Although this number appears low in comparison to work that applied the iNs system to the study of Alzheimer’s and which found >700 DEGs in iNs derived from participants with Alzheimer’s compared to controls, it is in line with a post-mortem study that conducted microarray-based transcriptome analysis on post-mortem brain samples derived from PLWH and PWOH (*11, 17*). In this post-mortem study, around 90 probes were significantly regulated in PLWH with no or only slight neurocognitive impairment compared with PWOH while analyzing gross brain tissue (i.e., not only neurons) across the neostriatum, neocortex, and white matter (*11*).

Interestingly, the authors found the majority of DEGs in the neostriatum (>80 regulated probes) and less than ten in the neocortex and white matter, respectively (*11*). Given the profound differences in numbers and genes of the different brain compartments, it would be interesting to apply different iNs protocols to our cohort of virologically suppressed PLWH and PWOH to identify putative DEGs in iNs of distinct neuronal subtypes in a future work (*19*).

We also observed increased expression of *SORCS1* in PLWH iNs. SORCS1 is a general regulator for intracellular trafficking, important for maintaining neuronal functionality, and it is associated with Alzheimer’s disease) (*29, 41*). Notably, the *SORCS1* gene encodes multiple protein isoforms (variant A, B, and C) with different trafficking and interacting properties (*29, 73*). Thus, to elucidate a putative role in HIV-1-related neurocognitive decline based on our observation, more detailed analyses with respect to these isoforms must be performed.

Finally, we observed increased expression levels of the inflammatory gene *IFI27* in iNs derived from PLWH compared to PWOH, which is in line with previously conducted post-mortem studies (*11, 69, 71*). Interestingly, this gene in particular has been associated with HIV-1 in several very recent studies (*74–77*). Mackelprang et al. found *IFI27* to become upregulated in blood samples of PLWH during acute infection and that it remained upregulated in the chronic state (*74*). The authors concluded that persistent elevation of a narrow set of interferon-stimulated genes including *IFI27* underlies chronic immune activation during HIV-1 infection (*74*). In this regard, our results suggest neuron-derived *IFI27* in this model as well. Liu et al. specifically searched for genes associated with immunological non-responders to HIV-1 infection in blood samples and found that *IFI27* expression levels are negatively correlated with CD4^+^ T cell count in PLWH (*76*). Moreover, they showed that the predictive power of *IFI27* expression levels in distinguishing PLWH with poor immune recovery was significant in their study and thus concluded that *IFI27* exhibits promising properties as biomarker for CD4^+^ T cell recovery. In our cohort, *IFI27* expression levels in PLWH iNs were likewise negatively associated with CD4^+^ T cell counts, which supports their findings in the context of neurons. In yet another study, Huang et al. recommended *IFI27* as a novel therapeutic target for HIV infection, which was based on differential expression analysis, and PPI network analysis using publicly available data sets on blood samples derived from PLWH and controls (*77*). Altogether, our findings in the context of PLWH-derived iNs support this recent association of *IFI27* with HIV-1 and suggest that its expression may also play a role in HIV-1-related neurocognitive impairment. Of note, in future studies, it would be interesting to investigate a potential link between the classification of NCI type I and NCI type II as introduced by Gelman et al. and the model of immunological non-responder following HIV-1 infection (*11*).

Besides *IFI27*, none of the other DEGs revealed here has been recognized in the few studies that compared differential genes expression in PLWH-and PWOH-derived post-mortem brain tissue samples (*11, 78, 79*). This is perhaps not surprising as our findings, focused on the comparison of iNs including a subset of glutamatergic and GABAergic neurons from PLWH and PWOH, have no direct comparator in the literature. In addition, we have performed unbiased whole-transcriptome bulk-RNA sequencing whereas prior studies on post-mortem brain tissue, e.g. those that identified *IFI27* as an upregulated gene in PLWH-derived samples, tended to conduct targeted gene expression profiling on a subset of previously selected genes (e.g., inflammatory genes) (*11, 69, 71*). Furthermore, our inclusion in this study of PLWH, carefully screened as per our methods to exclude those with significant neuropsychiatric confounds increases the sensitivity of our findings for HIV-1 related effects.

Concerning possible limitations of our study, we think that a comparison of six PLWH iNs samples to seven PWOH iNs has been sufficient to address the question whether gene expression of iNs differentiates virologically suppressed PLWH from PWOW. This is also consistent with previous studies on post-mortem brain samples in this context, which applied similar or even smaller cohort sizes (*11, 12, 79, 80*). Moreover, our two groups of PLWH and PWOH are very well-matched and the PLWH participants underwent extensive neurocognitive assessment and evaluation of HIV-1-related clinical parameters.

Taken together, we have been the first to study the effects of HIV-1 infection, and infectious disease in general, on neuronal gene expression using the model system of participant-derived iNs. We have found in our cohort that the resulting gene expression of iNs differentiates virologically suppressed PLWH from PWOH by identifying 29 DEGs between the two groups. From here on, subsequent studies should follow by focusing on the single genes identified by us while also including different neuronal subtype iNs protocols, and assays to assess neuronal functionality.

## Methods

### Study participants

The study was approved by the Rockefeller University Institutional Review Board. Written informed consent was obtained from all participants prior to their entering the study. Enrolled participants underwent a medical history, physical and neurological examination and psychiatric and substance use history at the screening visit. For PLWH and PWOH exclusion criteria adapted from Rippeth et al. (*81*) included severe neurological or diagnostic and Statistical Manual Fourth Edition-Text Revision (DSM-IV-TR) (*82*) psychiatric illness that affects cognitive functioning (e.g. schizophrenia, bipolar affective disorder), current diagnosis of major depressive disorder as assessed by the patient health questionnaire nine item depression scale (PHQ-9) (*83*) and not on stable antidepressant medication greater than 30 days, a history of head injury with loss of consciousness more than 30 min, DSM-IV-TR diagnostic criteria for alcohol or illicit substance abuse or dependence, not in remission, within 1 year of the screening visit (excluding marijuana), moderate or higher efavirenz attributable central nervous system (CNS)-related toxicity or serologic evidence of untreated syphilis or positive hepatitis C serology. All PLWH had documented treatment for at least 1 year with cART and plasma HIV-1 RNA levels below 50 copies/ml for a minimum of 6 months prior to study entry.

### Neuropsychological evaluation

The neuropsychological evaluation of the six PLWH recruited at the Rockefeller University was performed as described previously (*25*). Comprehensive neuropsychological evaluation assessing seven cognitive domains associated with HAND (attention/working memory; processing speed; learning; recall; abstraction/executive functioning; verbal fluency; and motor skills) was adapted from Rippeth et al. (*81*) and performed at the study visit by a study psychometrist. Using methods that correct for age, education, sex and ethnicity where appropriate, raw scores for all tests were transformed into T-scores (*81*). T-scores were then converted to deficit scores, which range from a minimum of 0 in the case of no impairment, to a maximum of 5 (*3, 84*). Calculating the sum of all deficit scores in the testing battery and then dividing by the number of administered tests allowed for determination of the global deficit score (GDS) for each participant, which provides a continuous measure of neurocognitive impairment (NCI).

### Dermal fibroblast isolation and propagation

Skin samples from all six PLWH and two of seven PWOH as detailed in the manuscript were collected via skin punch biopsy under Rockefeller University IRB-approved protocol. Dermal fibroblasts were isolated from skin biopsy samples and expanded by the MSK Stem Cell Research Facility. Briefly, a 6 mm diameter skin biopsy was dissected into 10 -15 smaller pieces, which were then plated on a 10 cm dish. Two to three pieces of samples were transferred into each well of a 6-well plate coated with 0.1 % gelatin and containing 500 µl fibroblast culture medium. The culture medium consisted of DMEM high glucose (ThermoFisher) supplemented with 10 % fetal bovine serum (HyClone), 1X NEAA (ThermoFisher), and 1X L-glutamine (ThermoFisher). A circular coverslip (FisherScientific) was carefully placed on top of the biopsy samples, and 1.5 ml of fibroblast culture medium was added onto the coverslip. Fibroblasts were observed approximately two weeks after plating and were passaged after three weeks onto gelatin-coated plates using trypsin-EDTA (0.05 % EDTA) for expansion. Dermal fibroblasts from five of seven PWOH control participants, chosen to match the demographics of the PLWH, were obtained via MTA from the *Coriell Institute for Medical Research* (NJ, USA).

### Generation of induced neurons

The performed protocol to generate induced neurons (iNs) was adapted from (*17, 24*). For the generation of lentiviral vectors, six million HEK293T cells were seeded into a 0.1 % gelatine-coated T150 cell culture flask in DMEM (Gibco) supplemented with 10 % fetal calf serum (FCS). The next day, cells were transfected with 6 µg of the packaging plasmid *psPAX2* (Addgene #12260), 6 µg of the envelope plasmid *pMD2.G* (Addgene #12259), and 6 µg the transfer plasmid *pLVX-UbC-rtTA-Ngn2:2A:Ascl1* (Addgene #127289) (*23*). The transfection mix containing polyethylenimine (60 µg/ml) (Polysciences) in 1 ml DMEM, and the three plasmids was added to the cells after an incubation period of 30 min. At 24 h post-transfection, the culture medium was exchanged to 15 ml fresh DMEM supplemented with 10 % FCS. At 48 h post-transfection, lentiviral vectors were harvested by pelleting any cells and cell debris at 400 x *g* for 5 min, aliquoting the resulting supernatant and storing at -80°C. UNA fibroblasts were generated by transducing 500,000 dermal fibroblasts with 500 µl of the lentiviral vector stock in a T25 cell culture flask while adding 4 µg/ml polybrene (Tocris) to improve transduction efficiency. At 24 h post-transduction, the medium was exchanged to fresh fibroblast medium. Selection with puromycin (1 µ/ml) (Sigma) started 72 h post-transduction. Transduced and selected fibroblasts (UNA fibroblasts) were passaged once and then frozen in liquid nitrogen. To generate iNs, all UNA fibroblasts were thawed the same day and processed in parallel to reduce batch effects during RNA sequencing. UNA fibroblasts were passaged several times after thawing and the neuronal conversion was performed as previously described (*24*). At day 21 of neuronal conversion, successfully converted iNs were isolated by FACS via staining for the neuronal marker PSA-NCAM. For this, cells were detached from the cell culture flasks with trypsin-EDTA and collected in FACS buffer consisting of 5 % FCS in PBS. Cells were pelleted (400 x *g*, 5 min) and incubated in a total of 200 µl FACS buffer containing the PE-conjugated PSA-NCAM antibody (Myltenyi Biotec) at a 1:100 dilution. After an incubation for 30 min at 4°C in the dark, the cells were washed twice with 500 µl FACS buffer and resuspended in 300 µl FACS buffer containing 1x DAPI (Thermo Scientific) used as live/dead stain. Cell sorting was performed on a BD FACSymphony™ S6 Cell Sorter at the WCM CLC Flow Cytometry Core Facility. Sorted cells were pelleted and then either lysed according to the respective downstream protocol or cultured in BrainPhys medium (StemCell) supplemented with B27 (1x) (Thermo), N2 (1 %) (StemCell), GDNF (20 ng/ml) (StemCell), BDNF (20 ng/ml) (StemCell), db-cAMP (500 µg/ml) (StemCell), and Laminin (1 µg/ml) (Thermo). For the first 24 hours post-FACS, we supplemented the medium with 10 µM ROCK inhibitor (MedChemExpress).

### Microscopic analysis and immunocytochemistry

Medium was removed, and the cells incubated in 4 % PFA for 20 min at room temperature (RT). After washing with PBS, cells were permeabilized with 0.1 % Triton-X-100 in PBS for 10 min at RT. After two additional washing steps with PBS, a blocking solution (2 % BSA in PBS) was applied for 1 h at RT. Primary antibodies were diluted in blocking solution and the cells were incubated with this antibody solution overnight at 4°C. Cells were washed twice with PBS and incubated with the secondary antibodies and Hoechst to stain nuclei for 2 hours at RT in the dark. Microscopic images were taken with the Olympus IX81 microscope (Olympus) using the *Slidebook* (version 6) software (3i). Image analysis has been performed with Fiji (*85*). The following antibodies were used in this study: Mouse-anti-Tubulin beta-3 (TUBB3) antibody (BioLegend), chicken-anti-MAP2 antibody (ThermoFisher), Alexa 488-conjugated goat anti-mouse IgG (ThermoFisher) and Alexa 568-conjugated anti-chicken IgG (ThermoFisher).

### Single-cell RNA sequencing and analysis

Cells were pelleted after FACS by centrifugation at 1,200 x *g* for 10 min. Medium was removed until only about 1 ml was left on top of the cells. The cells were resuspended and transferred into 1.5 ml tubes. After another centrifugation step (1,200 x *g* for 5 min), the complete supernatant was removed, and cells resuspended in 50 µl PBS containing 0.04 % BSA. Single-cell (sc)RNA sequencing has been performed at the Genomics Resources Core Facility (GRCF) at Weill Cornell Medicine. In brief, the 10X Libraries were sequenced on the Illumina NovaSeq6000 platform with pair-end reads (28 bp for read 1 and 90 bp for read 2). Sequencing data were analyzed by the 10X Cell Ranger pipeline (v7.1.0) in two steps. In the first step, Cell Ranger mkfastq demultiplexed samples and generated FASTQ files and in the second step, Cell Ranger count aligned FASTQ files to the 10X pre-built human reference genome (refdata-gex-GRCh38-2020-A) with standard parameters as described on 10X Genomics (https://www.10xgenomics.com/support/software/cell-ranger/latest/analysis/running-pipelines/cr-gex-count) and extracted gene expression UMI counts matrix. Count matrices were processed in RStudio using the *Seurat* package (*86–88*). Cells were filtered as previously described (>300/<10,000 unique feature counts and < 30 % mitochondrial reads), which resulted in 5994 (Participant 4642) and 8368 cells (Participant 3962) for downstream analysis (*17*). Data was normalized using the global-scaling normalization method with *LogNormalize* (scale factor 10,000). A subset of 2,000 features with high cell-to-cell variation was identified and scaled for downstream analysis. UMAP plots were generated using the identified dimensionality during principal component analysis. Percentages of cells expressing certain genes were determined using the *scCustomize* package (*89*).

### Bulk-RNA sequencing and analysis

Total RNA was extracted using the RNeasy kit (Qiagen) including the 15 min on-column DNase treatment. RNA integrity and quantity has been determined using the TapeStation instrument (Agilent). All RNA samples exhibited an RNA integrity number (RIN) above 9.0. Libraries were sequenced with paired-end 50 bps on a NovaSeqXplus sequencer. Raw sequencing reads in BCL format were processed through bcl2fastq 2.20 (Illumina) for FASTQ conversion and demultiplexing. After trimming the adaptors with cutadapt (1.18), RNA reads were aligned and mapped to the GRCh38 human reference genome by STAR (2.5.2) and transcriptome reconstruction was performed with Cufflinks (2.1.1) (*90, 91*). Raw read counts per gene were extracted using HTSeq-count v0.11.2 (*92*). Read count matrices were important into RStudio and differential gene expression analysis was performed using the *DESeq2* package (*93*). Low count genes (>10 reads) were pre-filtered and effect sizes were shrinked for visualization in MA plots using the *apeglm* method (*94*). Statistical significance was tested via the in *DESeq2* implemented Wald test (p-value) and corrected for false discovery rates (FDR) using the Benjamini-Hochberg method (p-adj.) DESeq2 median ratios count normalization was used. For visualization and cluster analysis, count data transformation was performed using the *vst* function (*95*). Gene set enrichment analysis was conducted with *EnrichR*, which uses Fisher’s exact test or the hypergeometric test (p-value) and FDR correction via the Benjamini-Hochberg method (p-adj.) (*96–98*). Gene sets used for this study were derived from Gene Ontology (*99, 100*), Jensen DISEASES (*64*), SynGO (*101*), Reactome (*102, 103*), and DisGeNet (*65*) databases.

### Protein-protein interaction (PPI) network analysis

We determined 1^st^-order interaction partners using the free open-source IntAct Molecular Interaction Database system (EMBL-EBI) (*62*), and the human reference interactome (HuRI) map (Center for Cancer Systems Biology at Dana-Farber Cancer Institute) (*63*). We generated and retrieved lists of the 1^st^-order interaction partners of the here identified DEGs based on the underlying literature curation and direct user submission (IntAct) as well as the un-biased, systematic, yeast two-hybrid screen for PPIs (HuRI).

### Statistics and software

Besides the beforementioned software, *GraphPad* was used for a subset of statistical analysis and plots. *Inkscape* was used for illustrations and finalization of figures. *BioRender* was used to generate a subset of schemes. Statistical analyses were run with either *RStudio* using the *DESeq2* or *Seurat* package, *GraphPad*, or *EnrichR* and has been depicted throughout the manuscript where applied.

## Data availability

Please contact the corresponding author Teresa H. Evering for any inquiries regarding the used material and uploaded data. Raw data derived from the bulk-RNAseq and scRNAseq experiment will be made publicly available through the Gene Expression Omnibus genomics data repository (https://www.ncbi.nlm.nih.gov/geo/) upon publication. Complete lists of differentially expressed genes are available as supplementary material.

## Acknowledgements

Many thanks to Fred Gage, Jerome Mertens, and especially Larissa Traxler for their support and help in setting up the iN protocol. We want to thank all members of the Nixon and Furler group for thoughtful discussions and help. We thank Aaron Zhong for helping with the lab work and Johannes Ptok for helping with the bioinformatic analysis. We thank Anand Ramani for sharing his expertise in neuronal cell culture. We thank the members of the WCM CLC Flow Cytometry Core Facility for their services and help in fluorescence-activated cell sorting of the iNs. We thank the Genomics Resources Core Facility (GRCF) at Weill Cornell Medicine for their services and great help regarding the transcriptomic analyses.

## Author contributions

Conceptualization: THE

Methodology: PNO, YW, MS, THE

Investigation: PNO, YW, SB, MS, MAS, LSTB, SM, AH, THE

Visualization: PNO, SB, AH

Funding acquisition: TZ, THE

Project administration: TZ, THE

Supervision: RBJ, DFN, MY, LCN, TZ, THE

Writing – original draft: PNO

Writing – review & editing: All authors

## Data and materials availability

Please contact the corresponding author Teresa H. Evering for any inquiries regarding the used material and uploaded data. Raw data derived from the bulk-RNAseq and scRNAseq experiment will be made publicly available through the Gene Expression Omnibus genomics data repository (https://www.ncbi.nlm.nih.gov/geo/) upon acceptance of the article. Complete lists of differentially expressed genes between our groups are available as supplementary material.

---

**Fig. S1.**
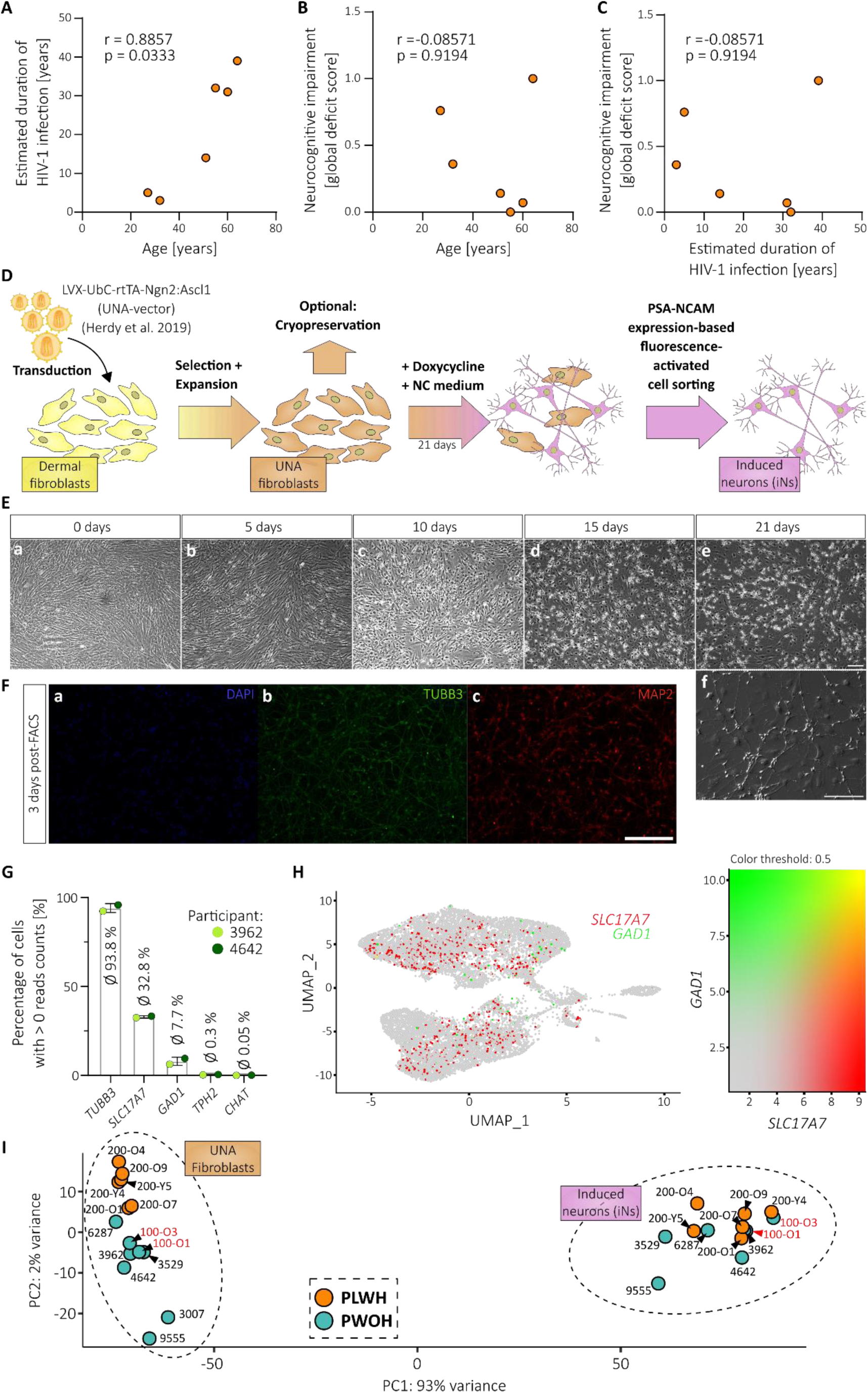
Study cohort and induced neuron (iNs) characteristics. **(A-C)** Correlation of age, estimated duration of HIV-1 infection, and neurocognitive impairment in the PLWH study group (n =6). Data points are individual values and correlation analysis performed via two-tailed, nonparametric spearman. **(D)** Scheme illustrating the workflow of participant-derived iNs generation following the previously published *Mertens* protocol (*24*). **(E)** Microscopic images of participant-derived skin fibroblasts at different days during the transdifferentiation protocol. **(F)** Single channel microscopic images after immunocytochemistry of induced neurons (iNs) 3-days post-FACS stained for TUBB3 (TUJ1), MAP2 and nuclei (DAPI). **(E-F)** Scale bars are 20 µm. **(G)** Percentage of cells from the here performed scRNA analysis that express the annotated pan-neuronal and neuronal subtype marker genes. Data presented as individual data points with mean ± SD. **(H)** UMAP plot showing *SLC17A7* and *GAD1* expression patterns and values among iNs. **(I)** PCA plot showing the calculated distance between bulk-RNA samples with annotations for every single study participant.

**Fig. S2.**
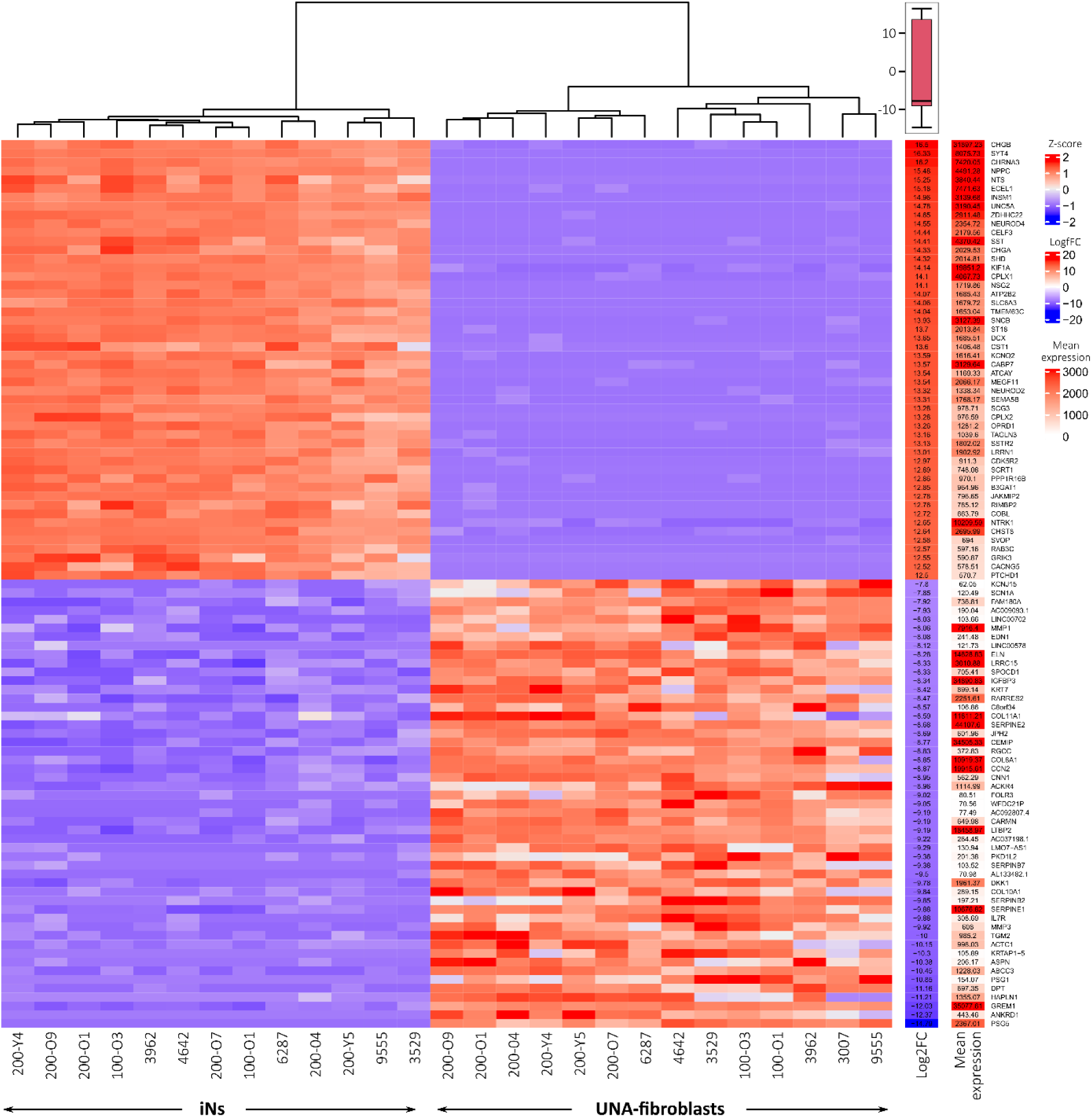
Heatmap showing clustering of UNA fibroblasts and iNs according to the top 50 up-and downregulated genes. This heatmap displays the mean expression values of the top 50 up-and downregulated statistically significant (p-adj. < 0.05, log2fc > +/- 0.5) DEGs and the resulting log2 fold change between the UNA fibroblast and iNs samples based on our bulk-RNA analysis.

**Fig. S3.**
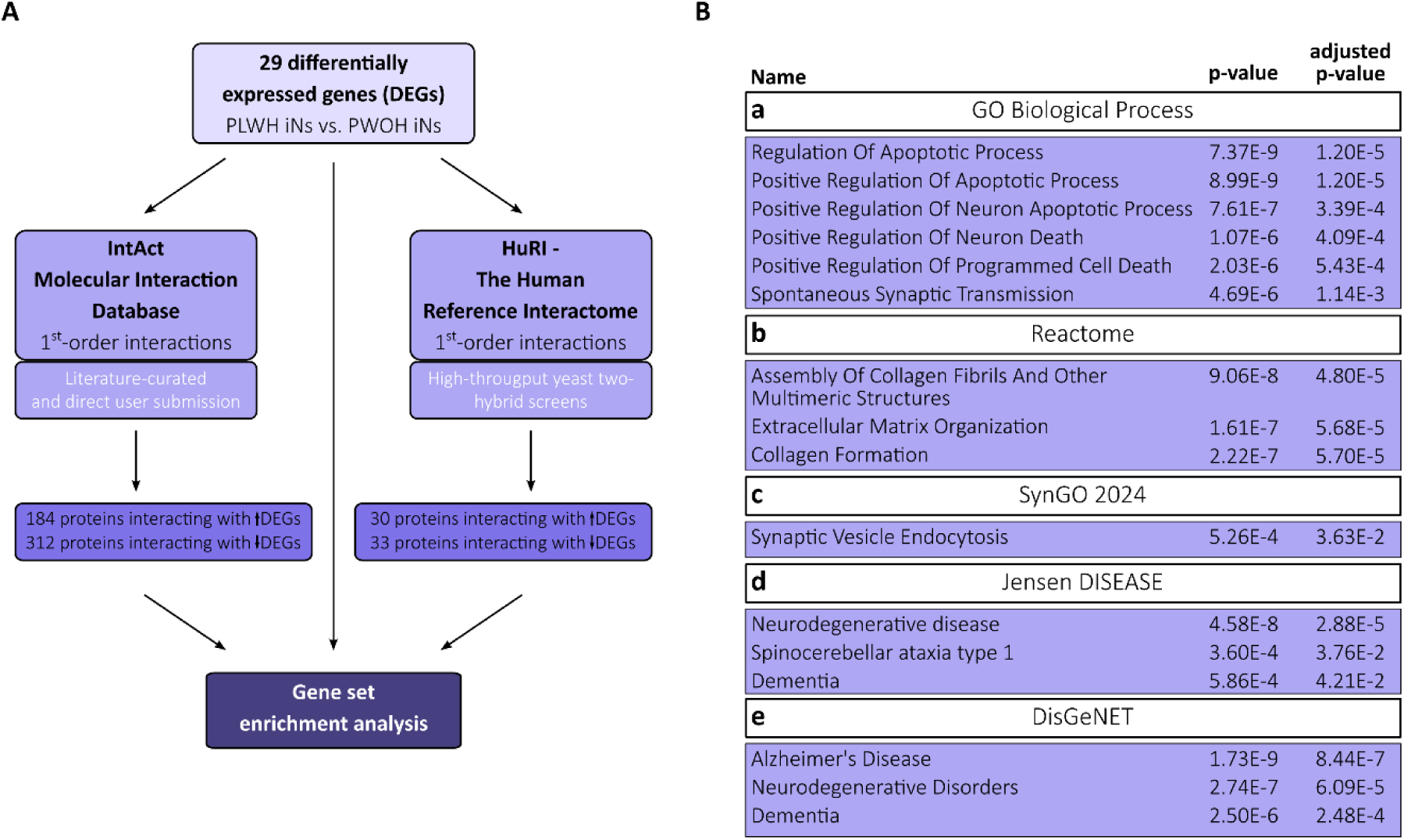
Protein-protein interaction network mapping supports differential ECM organization, and synaptic transmission in PLWH iNs and indicates neuronal apoptosis as another affected pathway. **(A)** Flowchart summarizing the performed protein-protein interaction (PPI) network mapping approach using the IntAct Molecular Interaction Database (*62*) and The Human Reference Interactome (HuRI) (*63*). **(B)** Gene set enrichment analysis-derived terms significantly enriched within the obtained PPI network of 1^st^-order interaction partners of the here identified 29 DEGs between PLWH-and PWOH-derived iNs.

**Fig. S4.**
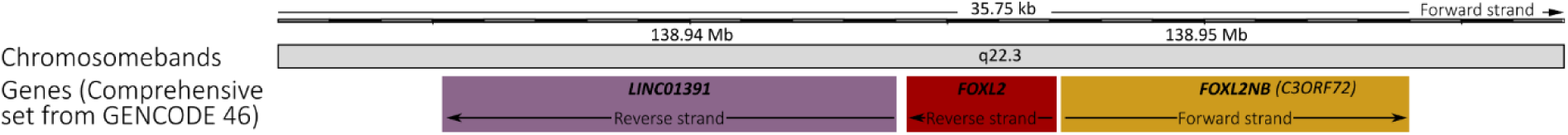
Gene locus of *LINC01391*, *FOXL2*, and *FOXL2NB*. The exact location of the three genes *LINC01391*, *FOXL2*, and *FOXL2NB* on human chromosome 3 is shown. Information and original graphic retrieved from the freely available genomic resource Ensembl (https://www.ensembl.org; 9^th^ June 2024, 12:33) (*104*).

**Fig. S5.**
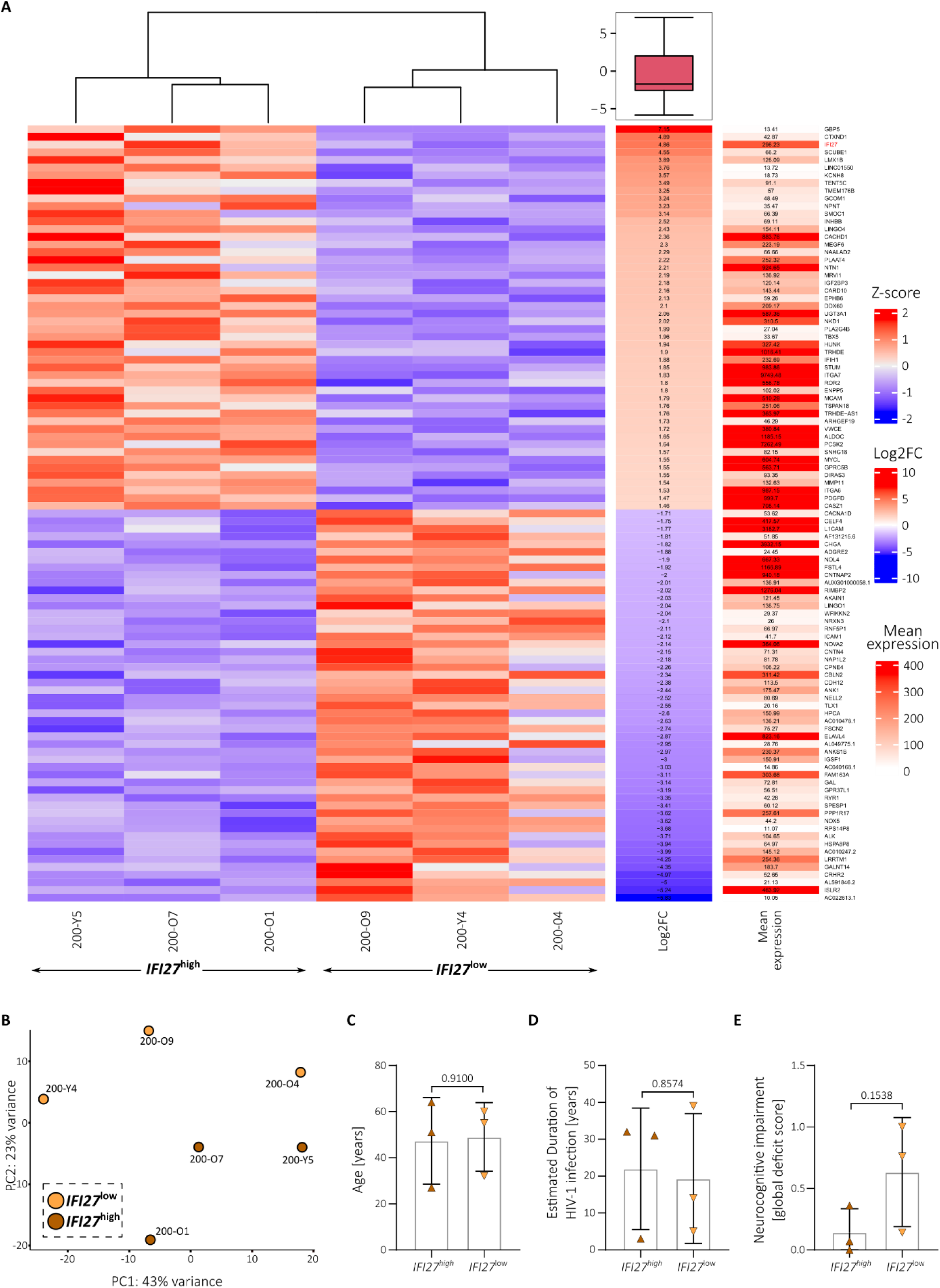
Genes associated with and biological parameters of the *IFI27*^high^ and *IFI27*^low^ expressing PLWH participants. **(A)** Heatmap showing the mean expression values of the top 50 up-and downregulated statistically significant (p-adj. < 0.05, log2fc > +/- 0.5) DEGs and the resulting log2 fold change between *IFI27*^high^ and *IFI27*^low^ expressing PLWH iNs based on our bulk-RNA analysis. (B) PCA plot showing the calculated distance between the *IFI27*^high^ and *IFI27*^low^ expressing PLWH iNs samples. **(C-E)** Age **(C)**, estimated duration of HIV-1 infection **(D)**, and neurocognitive impairment measured as global deficit score **(E)** of PLWH (n = 6) divided into *IFI27*^high^ vs. *IFI27*^low^ expressing participants. **(A, D, G)** Statistical significance tested with unpaired, two-tailed *t*-test. Data presented as individual data points with mean ± SD and p-value.

